# Adaptive immune responses are not causal to SGN death after kanamycin-induced hair cell loss

**DOI:** 10.64898/2026.07.24.740643

**Authors:** Adrianna M. Caro, Steven H. Green

**Affiliations:** Department of Biology, University of Iowa, Iowa City, IA 52242

## Abstract

Aminoglycoside antibiotics such as kanamycin induce sensorineural hearing loss by killing hair cells, resulting in secondary degeneration of spiral ganglion neurons (SGNs). Previous studies show that anti-inflammatory agents reduce SGN death, implicating a causal role of the immune response. This is consistent with observations of increased numbers of macrophages and lymphocytes, including T and NK cells, in the spiral ganglion after exposure to aminoglycosides. Here, we directly test the role of T cells and other lymphocytes in SGN degeneration in kanamycin-deafened rats. Homozygous RNU nude rats that lack T cells – but retain NK and B cells– show neurodegeneration similar to rats with a normal T cell complement, indicating that T cells are not necessary for neurodegeneration. Homozygous SRG rats lacking all lymphocytes (i.e., T, B, and NK cell-deficient), exhibit remarkable regional variation in the pattern of spiral ganglion degeneration post-deafening. In the basal half of the ganglion, SGN degeneration is significantly reduced in deafened SRG rats, implying a role for lymphocytes, presumably NK cells of the innate immune system, in SGN death. In the apical half of the deafened ganglion, SGN degeneration is not significantly affected by the lack of all lymphocytes, implying a role for other cellular mechanisms.

## Introduction

Sensorineural hearing loss is typically caused by damage to cochlear hair cells, the auditory sensory receptors, or to their afferent spiral ganglion neurons (SGNs). Hair cells are highly susceptible to damage from a variety of insults, and their loss can result in subsequent death and degeneration of SGNs. This neurodegeneration has been investigated in several animal models of hair cell loss caused by aminoglycoside ototoxicity, including in rats (Alam et al., 2007; Kopelovich et al., 2013), guinea pigs (Landry et al., 2011; Miller et al., 2007), and cats (Leake et al., 2011; Leake et al., 2013). The rate of SGN death is species specific, but is generally slow, occurring over a period of months in rats and guinea pigs, to years in cats. In the rat, over 80% of SGNs are lost over the course of ∼14 weeks following aminoglycoside-induced hair cell loss (Alam et al., 2007).

Accumulating evidence implicates an inflammatory/immune response as a cause of SGN death (Rahman et al., 2023; Shimada et al., 2023). Macrophages and lymphocytes, including CD4+ T cells, CD8+ T cells, and CD161+ NK cells increase in number in the spiral ganglion after deafening due to hair cell loss, with the increase in macrophages underway even prior to the onset of SGN death (Caro et al., 2026). Suppressing this response with anti-inflammatory agents significantly reduces SGN death, implying that this response is causal, at least in part, to SGN death post-deafening (Rahman et al., 2023). A complicating factor is that there are regional and temporal differences within the ganglion in the expression of pro-inflammatory cytokines (Rahman et al., 2023), in the number of macrophages, and in the rate of SGN death (Caro et al., 2026). Additionally, an immune response is itself complex, consisting of innate and adaptive immune responses, mediated by macrophages and lymphocytes that communicate bi-directionally.

T cells are central regulators of the immune response, being activated by MHCII-expressing macrophages to initiate an adaptive immune response (Roche & Furuta, 2015). T cells reciprocally modulate macrophage and microglial activation states through cytokines such as IFN-γ, IL-17, and IL-10 (S. Chen et al., 2023; Reel et al., 2024). The presence of MHCII-expressing macrophages, as well as CD4+ helper T cells (Caro et al., 2026; Gansemer et al., 2024) in the spiral ganglion, raises the possibility that, after hair cell loss, macrophages activate an adaptive immune response that is causal to SGN death. Nevertheless, the presence of NK cells in addition to macrophages (Caro et al., 2026) might imply that an innate immune response is sufficient to cause SGN death.

To assess the role of T cells and other lymphocytes in spiral ganglion neurodegeneration, we used a well-established model of deafening – destruction of hair cells by aminoglycosides in the rat (Alam et al., 2007; Bailey & Green, 2014; Gansemer et al., 2024; Rahman et al., 2023) – together with genetic manipulations to selectively delete T cells or delete all lymphocytes. These results show that T cells are not necessary for SGN death post-deafening, arguing against a causal role for the adaptive immune system. The effect of deleting all lymphocytes, including innate NK cells, on SGN death differs by location in the spiral ganglion, revealing regional differences in neuroimmune interactions and neurodegeneration.

## Materials and Methods

### Animals and aminoglycoside deafening procedure

For all experiments, male and female rats were used. The day of birth was assigned as postnatal day 0 (P0) and rats were euthanized at P70. Rats were housed under a 12-hour light/dark cycle and were allowed to feed ad libitum throughout the experimental period. Protocols and procedures for all animal experiments were approved by the University of Iowa Institutional Animal Care and Use Committee.

All rats were deafened neonatally by daily intraperitoneal injections of kanamycin sulfate (400mg/kg) from P8 to P16. As previously reported, this protocol results in the loss of outer hair cells by the end of kanamycin treatment, while inner hair cell loss is complete by approximately P20 (Bailey & Green, 2014). Deafening was verified by assessment of auditory brainstem response (ABR) and/or by lack of remaining hair cells detected by immunofluorescence. Any animals with surviving hair cells were excluded from the study.

Adult RNU rats were purchased from Charles River to form our breeding colony. RNU heterozygous (Foxn1^rnu/+^) females were bred to RNU homozygous (Foxn1^rnu/rnu^) males to produce litters of heterozygous and homozygous pups. RNU heterozygotes (“hets”) have a normal T cell population, equivalent to wild-type rats, while RNU homozygotes (“RNU nudes”) lack a functional thymus and therefore are depleted of functional T cells (Rolstad, 2001). RNU “nudes” also lack a coat, so can be readily distinguished from heterozygous littermates, which have a normal coat.

Pregnant SRG dams and adult male mice were purchased from Charles River. SRG rats are of a Sprague Dawley background and are Rag2^-/-^, Il2rg^-/-^ resulting in a lack of mature T, B, and NK cells. Because of their severe immunodeficiency, SRG rats were housed in a barrier facility until the day prior to ABR testing and euthanasia. Sprague-Dawley rats from our breeding colony or purchased from Envigo were used as wild-type (WT) controls for SRG experiments.

### Auditory brainstem response (ABR) test

ABR recordings were measured between P55-P69 to assess auditory threshold in immunodeficient rats. As previously described (Hu, 2020, Gansemer, 2024; Hemachandran, 2024), rats were anesthetized with ketamine/xylazine and an active needle electrode was placed at the midline of the vertex of the skull, a reference electrode at the ipsilateral mastoid, and a ground electrode in the lower back. Tone pip stimuli were delivered to the external auditory meatus via a custom-made insertion tube connected to an MF1 speaker (Tucker-Davis Technologies, Inc.). Tone pip stimuli of 5 ms duration, presented at a rate of 21/s, alternating polarities, were presented at 4, 8, and 16 kHz (RNU rats) or 8, 16, and 32 kHz (SRG and PLX-treated rats). Responses were recorded from 90 dB SPL to 10 dB below threshold in 10 dB steps and repeated in 5 dB steps near threshold. The lowest stimulus level at which there was a detectable and repeatable wave I was considered threshold. The wave I amplitude was measured between the first positive maximum to the following negative minimum. Wave I latency was defined as the time, in ms, from the start of the stimulus (0 ms) to the wave I peak.

Tests were performed using an RZ6 multi I/O processor with a RA4PA 4 channel preamplifier (Tucker-Davis Technologies, Inc.) in a custom-made sound-proof chamber. BioSigRZ (Tucker Davis Technologies, Inc.) was used to generate and deliver the stimuli and record response signals. A high-pass filter was set at 3000 Hz and a low-pass filter at 300 Hz. Final signals are an average of ∼512 sweeps. ABR data were exported from BioSigRZ and off-line processing and analysis were performed using custom-written scripts in MATLAB. The MATLAB scripts used for processing and analysis are available at https://github.com/bgansemer/ABR-analysis.

### Cochlear histology preparation

One cochlea chosen at random from each rat was used for histological analysis of hair cells, neurons, and leukocytes. Rats were deeply anesthetized with ketamine/xylazine and transcardially perfused with ice-cold PBS followed by ice-cold 4% paraformaldehyde in 0.2 M phosphate buffer. Spleens were harvested from some animals for use as positive controls for immunodetection of leukocyte markers. Cochleae were dissected from temporal bones and fixed in 4% paraformaldehyde at 4°C for up to 24 hours, followed by decalcification for 4-10 days in 0.12M EDTA, with EDTA refreshed approximately every 3 days until decalcification was complete. Cochleae were cryo-protected by sequential 30-minute incubations in 10%, 15%, 20%, and 25% sucrose, then overnight in 30% sucrose in PBS. Cochleae were then transferred to O.C.T. compound (Tissue-Tek) and maintained on a rotator overnight, then embedded in O.C.T. compound, oriented for sectioning, and flash-frozen in liquid nitrogen. Cochleae were stored at -80°C until sectioning. Serial sections (25 μm in thickness) were cut parallel to the mid-modiolar (central) plane on a Leica Cryostat and collected on Fisher Superfrost slides. Sections were stored at -20°C prior to immunolabeling.

### Immunofluorescence

A minimum of three mid-modiolar sections separated by 50 μm (i.e., every third section) were used for immunofluorescence imaging and cell counting. Immunofluorescence labeling was performed as described in Caro et al. (2026). Briefly, cochlear sections were permeabilized with 0.5% Triton X-100 in Tris-buffered saline pH 7.6 (TBS) for 20 minutes, then covered with blocking buffer (5% goat serum, 2% bovine serum albumin, and 0.02% sodium azide in TBS). After blocking for 3-4 hours at room temperature, sections were incubated in primary antibody in blocking buffer overnight at 4°C. After washing with TBS, sections were then incubated in secondary antibodies in blocking buffer for 3-4 hours at room temperature. Nuclei were stained with Hoechst 33342 (10 μg/ml in PBS, Sigma). Slides were coverslipped with Fluoro-Gel mounting medium in Tris-buffer, sealed with nail polish, and stored at 4°C for up to 96 hours until imaging.

Neurons were identified by labeling with antibody to neuron-specific β_III_-tubulin (TUJ1). Macrophages were identified by labeling with IBA1 antibody. Lymphocytes were identified as spherical cells, 8-15 µm in diameter with high nuclear to cytoplasmic ratio and as CD45+/IBA1-in sections labeled with both CD45 and IBA1 antibodies. Hair cell ablation was verified by lack of myosin 6/myosin 7a positive cells in the organ of Corti.

The following antibodies were used: anti-TUJ1 (Biolegend; cat. #80122, RRID AB_2313773); anti-IBA1 (Wako; cat. #019-19741, RRID AB_839504); anti-CD45 (Biolegend; cat. #202201, RRID AB_314003); anti-CD161 (Bio-Rad; cat. # MCA1427GA, RRID AB_566557); anti-CD68 (Abcam; cat. #31360, RRID AB_1141557); anti-myosin 6 (Sigma; cat. #M5187, RRID AB_260563); anti-myosin 7a (Proteus; cat. #25-6790, RRID AB_10015251). The following secondary antibodies were purchased from Thermo Fisher Scientific: Alexa Fluor 488 (cat. #A11008, RRID AB_143165); Alexa Fluor 546 (cat. #A21123, RRID AB_2535765); Alexa Fluor 633 (cat. #A21136, RRID AB_2535775).

### Imaging and quantitation

Imaging and quantitation were performed as we have previously described in Caro et al. (2026). Labeled sections were imaged on a Leica SPE confocal using a 40X (0.95 NA) or 63x (1.4 NA) objective, 1x digital zoom, and z-axis increments of 1 μm. Fiji/ImageJ (NIH) was used for all image analysis. All image stacks were assigned a random 8-digit code number using a proprietary Fiji plugin to blind the counter from the experimental condition. Cell counts were done on maximum intensity z-projections of the image stacks. Only cells with a visible nucleus were counted.

Cochleae were sectioned parallel to the mid-modiolar (central) plane. In each section, we assessed the spiral ganglion, which is contained within Rosenthal’s canal, in four distinct regions (from base to apex): the basal turn (termed “base”), the basal part of the middle turns (“mid 1”), the apical part of the middle turns (“mid 2”) and the apical turn (“apex”), as previously described (Caro et al., 2026). The former two are collectively referred to as “basal half,” and the latter two as “apical half.” Because of the spiral anatomy of the cochlea, in sections at or close to the mid-modiolar plane at least three turns, i.e., three cross-sections, of the spiral ganglion are visible in each section. The outline of Rosenthal’s canal in each turn was traced and its cross-sectional area measured. The cell count in each cross-section of Rosenthal’s canal was divided by the cross-sectional area to calculate the density.

### Flow Cytometry

Spleens were collected from control RNU het and RNU nude rats perfused with ice-cold PBS. Spleens were minced with a razor blade, immersed in Accutase (Innovative Cell Technologies, cat. #AT104) and incubated at 37°C for 10 minutes to liberate cells. Tissue was then forced through a 70-um strainer to generate a single cell suspension. Red blood cells were lysed with ACK lysis buffer. Single cell-suspensions were washed in FACS buffer (BD Biosciences, cat. #563503), centrifuged at 300 x g for 5 min at 4°C, and immediately labeled with an antibody cocktail for 30 min at 4°C containing the following antibodies: CD45-Pacific Blue (BioLegend; cat. #202225, RRID:AB_2721618), CD3-PE (BioLegend; cat. #201411, RRID:AB_2563272), and CD161-FITC (BioRad; cat. #MCA1427F, RRID:AB_321596). Cells were then centrifuged, washed in FACS, and resuspended in Cytofix/Cytoperm (BD Biosciences, cat. #554722) for 15 min at 4°C to fix the cells. Labeled cells were run through a BD LSR II cytometer and data were analyzed in FlowJo.

### Statistical analysis

Statistical analyses for all data were performed using GraphPad Prism, with specific tests used summarized in the figure legends. A Shapiro-Wilk normality test was performed for all data sets prior to testing for significance of differences. A p-value <0.05 was considered statistically significant.

## Results

### Foxn1^rnu/rnu^ (RNU nude) rats lack T cells

Athymic Foxn1^rnu/rnu^ nude rats lack mature T cells (Cash et al., 1993; Rolstad, 2001). We confirmed this by flow cytometry of the spleen (Figure 1), finding only ∼1.5% cells positive for the T cell marker CD3 in Foxn1^rnu/rnu^ rats, significantly reduced from ∼29% found in spleens from control Foxn1^rnu/+^ heterozygotes (Figure 1C). We could therefore use RNU rats to test whether T cells are necessary for SGN degeneration after hair cell loss. To this end, we compared SGN survival post-deafening between Foxn1^rnu/+^ RNU heterozygotes (RNU het) and T cell deficient Foxn1^rnu/rnu^ RNU homozygous mutant (RNU nude) rats.

**Figure 1.**
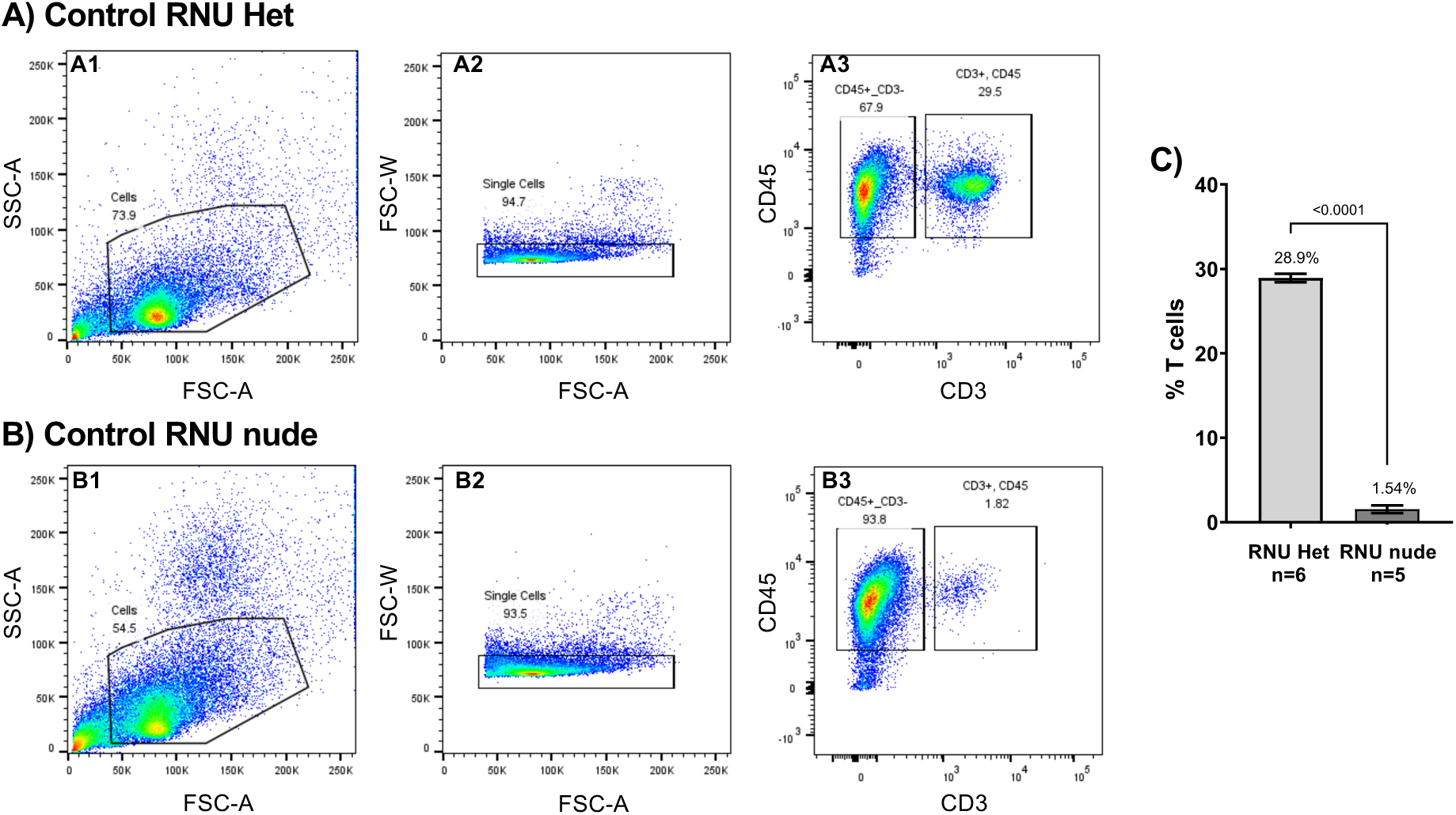
Foxn1^rnu/rnu^ nude rats are deficient of T cells. Representative plots from flow cytometric analysis of spleen from a **(A)** RNU heterozygote and **(B)** RNU nude rat. Panels show the gating strategy used to confirm depletion of CD3+ T cells in RNU nudes. **(A1, B1)** The first gate was set from the forward-scatter versus side-scatter to capture cells in the expected size and granularity range for lymphocytes. **(A2, B2)** Single cells were selected for by size; doublet cells were excluded. **(A3, B3)** Single cells were then plotted to evaluate expression of CD45 and CD3. Labels inside each plot indicate the target population captured within the gate. The numbers in each plot indicate the percent of cells represented within the gate. **(C)** Comparisons between RNU het and RNU nude rats indicate significant depletion of CD3+ T cells in RNU nudes. Shown are means ± SEM. n=number of animals. Significance determined by unpaired t-tests.

### T cell ablation in RNU nude rats does not affect normal SGN number nor SGN death after hair cell loss

After destroying hair cells with neonatal kanamycin injections as described in methods, we quantified SGN density in postnatal day 70 (P70) RNU rats at four cochlear locations along the base to apex axis. As described in (Caro et al., 2026), these locations are termed base and mid 1, collectively referred to as the basal half, and the mid 2 and apex, collectively referred to as the apical half. Representative images for each location are shown in Figure 2. We found no significant difference in neuron number between undeafened control RNU hets and undeafened control RNU nudes, indicating that the Foxn1 mutation itself does not affect SGN number in control rats.

**Figure 2.**
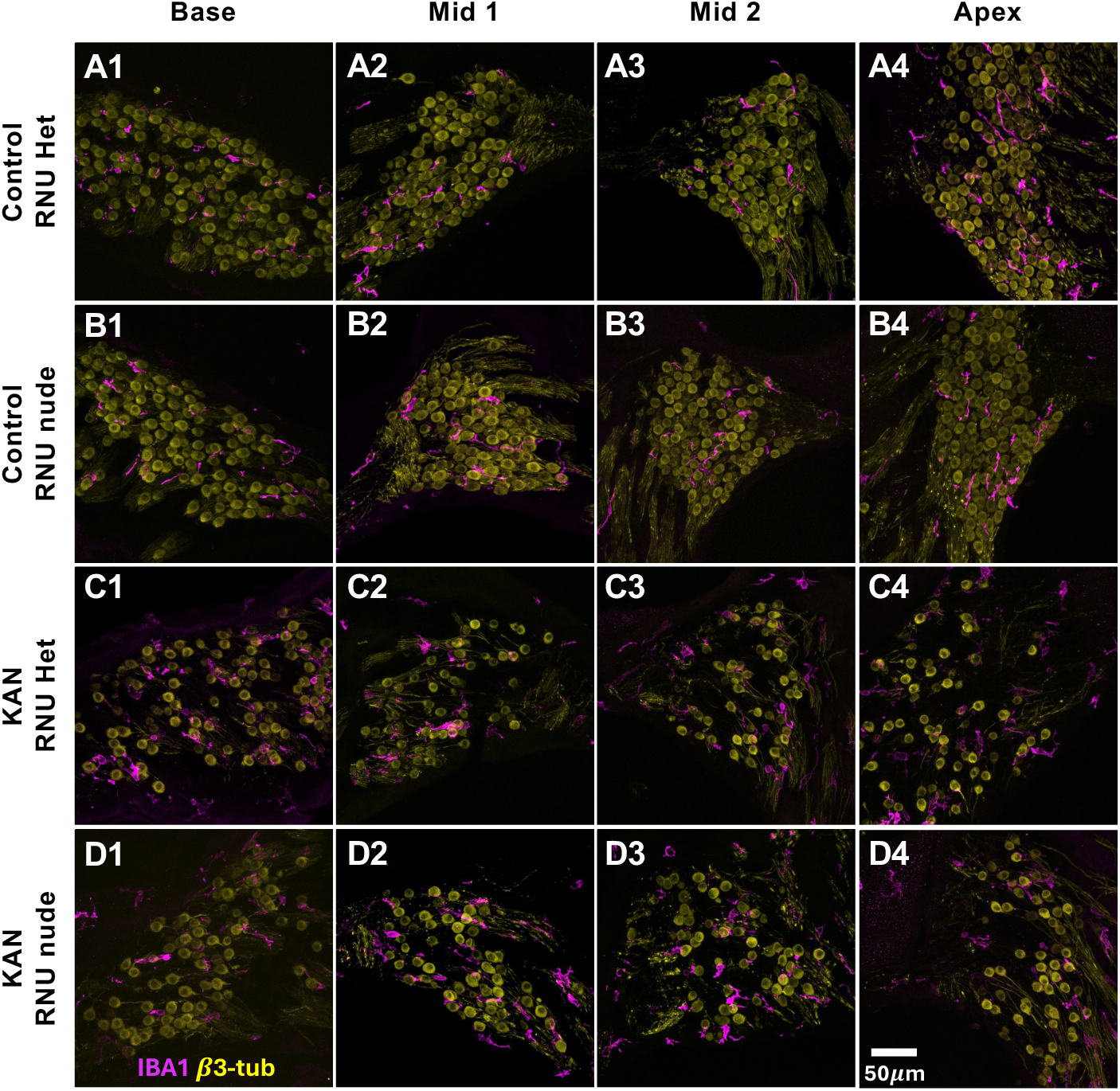
SGNs and macrophages during neurodegeneration in control and kanamycin-treated RNU rats. Shown are SGNs (βIII-tubulin, yellow) and macrophages (IBA1, magenta) in control T cell competent RNU heterozygotes (RNU het, **A1-A4**), control T cell deficient RNU nude (**B1-B4**), kanamycin treated (KAN) RNU het (**C1-C4**), and KAN RNU nude rats (**D1-4**) at postnatal day 70 (P70). Representative images from cochlear cross-sections at four different locations in the spiral ganglion along the base-to-apex axis (tonotopic) axis are in columns from left to right: column 1 corresponds to the base (column 1), mid 1 (column 2), mid 2 (column 3), and apex (column 4).

We found that, by P70, all kanamycin-treated (KAN) rats – both RNU hets and RNU nudes – had significantly reduced SGN numbers at all regions of the spiral ganglion relative to undeafened rats (Figure 3A). SGN counts showed no significant difference in SGN number between deafened RNU nudes or RNU hets in any region of the cochlea (Figure 3A). The lack of significant difference in the extent of SGN death between KAN RNU hets and T cell-deficient KAN RNU nudes implies that T cells are not required for SGN death after hair cell loss.

**Figure 3.**
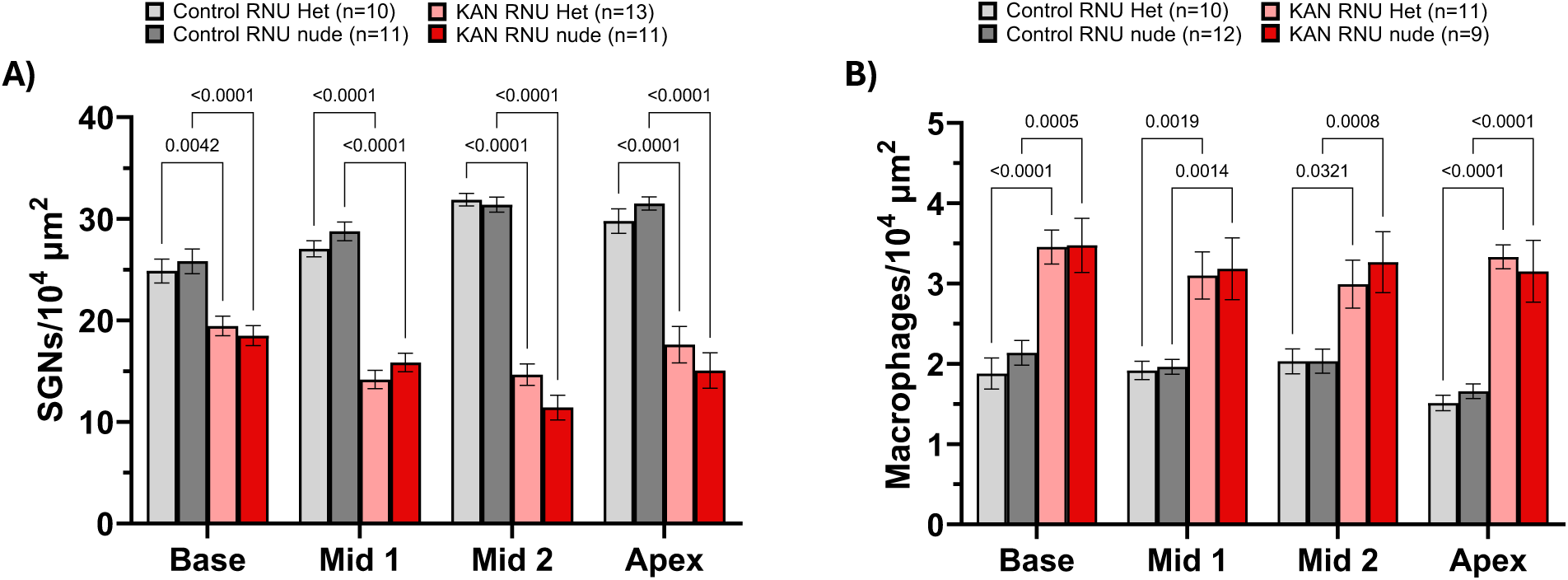
Quantification of SGNs and macrophages by cochlear location in control and KAN RNU rats at P70 indicate that T cells are not required for SGN death after hair cell loss. Number of (**A**) SGNs (βIII-tubulin+ cells) and (**B**) macrophages (IBA1+ cells) per 10^4^ µm^2^ at four locations in the spiral ganglion. Shown are means ± SEM. Significance of differences was determined using a two-way ANOVA with Tukey’s multiple comparisons.

### Macrophage number in the spiral ganglion increases similarly in KAN RNU hets and KAN RNU nudes after hair cell loss

We have previously shown that after hair cell loss, the number of macrophages in the spiral ganglion is significantly increased relative to controls (Caro et al., 2026; Gansemer et al., 2024; Rahman et al., 2023). While an important function of macrophages is recruitment and activation of T cells, T cells reciprocally modulate macrophage and microglial activation states (S. Chen et al., 2023; Reel et al., 2024). Thus, we asked whether T cells play a role in recruiting monocytes into the spiral ganglion of KAN treated rats. To that end, we asked whether lack of T cells in RNU nude rats had any effect on spiral ganglion macrophage number. We found that in control undeafened rats, there was no significant difference in macrophage number between control RNU het and control RNU nude rats (Figure 3B), indicating that the Foxn1 mutation does not affect the normal population of resident macrophages in the ganglion. In both undeafened control hets and control RNU nudes, macrophage density at all locations was approximately 2 cells/10^4^ *μ*m^2^. Similarly, there was no significant difference in macrophage number between KAN hets and KAN RNU nude rats. Macrophage density in both KAN hets and KAN RNU nude rats was approximately 3-3.5 cells/10^4^ *μ*m^2^ across all regions of the spiral ganglion, consistent with the number reported in deafened Sprague-Dawley wild-type rats (Caro et al., 2026) and significantly increased compared to undeafened controls (Figure 3B). These data indicate that T cells are dispensable for both the decrease in SGN number and the increase in macrophage number post-deafening.

### Lymphocytes are present in the spiral ganglia of both KAN RNU hets and KAN RNU nudes

RNU nude rats are expected to lack T cells but retain other types of lymphocytes. We indeed found that cells identified as lymphocytes (criteria are CD45+/IBA1-spherical cells, 8-15 µm in diameter, with a high nuclear to cytoplasmic ratio) were present in both KAN RNU hets (Figure 4A, C) and KAN RNU nude rats (Figure 4B, C). Representative low magnification images from the Mid 1 region are shown for control (Figure 4A) and KAN (Figure 4B) RNU nude rats.

**Figure 4.**
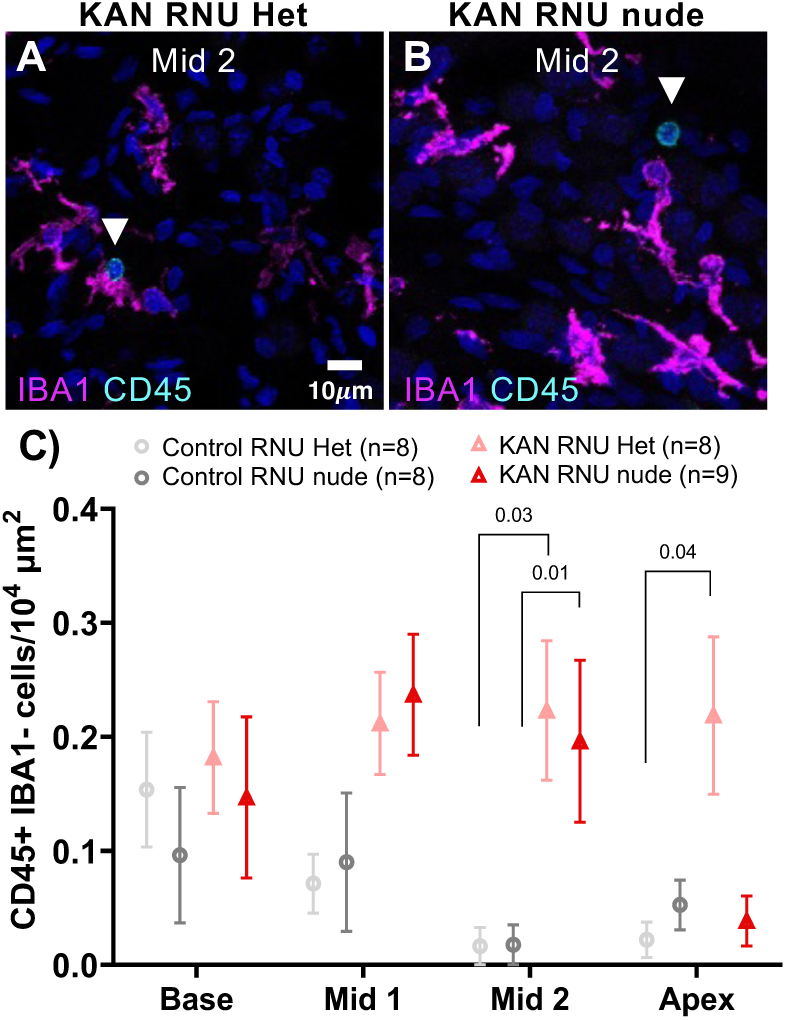
The number of CD45+/IBA1-cells increases in the spiral ganglion of KAN RNU het and KAN RNU nude rats. Representative high magnification images of spiral ganglia from the Mid 2 region of a **(A)** KAN RNU het and **(B)** KAN RNU nude rat labeled to show presence of CD45+/IBA1-leukocytes in the ganglion, emphasized by white arrows. These images show labeling with CD45 (cyan), IBA1 (magenta), and nuclei (blue, Hoechst 33342). Dotted outlines delineate Rosenthal’s canal. **(C)** Quantification of CD45+/IBA1-cells per 10^4^ µm^2^ at four locations in the spiral ganglion. Shown are means ± SEM. n=number of animals. Significance determined by one-way ANOVA Kruskal-Wallis test with Dunn’s multiple comparisons.

We have previously shown that lymphocytes, including CD161+ NK cells, CD4+ helper T cells, and CD8+ cytotoxic T cells, increase in number in the spiral ganglion after deafening in wild-type Sprague-Dawley rats (Caro et al., 2026). We asked whether the increase in lymphocytes within the ganglion persists in KAN RNU nude rats, despite their lack of T cells. We evaluated the number of remaining lymphocytes within the spiral ganglion of undeafened control and KAN RNU het and RNU nude rats. We found that in control RNU nude and RNU hets, a few lymphocytes were present within ganglia from undeafened rats (Figure 4C). In control undeafened rats, we found 0-0.15 lymphocytes/10^4^ *μ*m^2^ at all regions of the ganglion, consistent with our previous report (Caro et al., 2026). In deafened rats, both KAN RNU hets and KAN RNU nudes, lymphocyte density increased to 0.15-0.25 cells/10^4^ *μ*m^2^ in the base, mid 1, and mid 2 regions (Figure 4C), an increase significant only in the mid 2 region. In the apex, lymphocyte density was significantly increased for KAN RNU hets relative to undeafened control RNU hets, but, for KAN RNU nudes, lymphocyte density did not increase significantly post-deafening.

Given that RNU nude rats lack T cells (Figure 1C), we expected any CD45+/IBA1-lymphocytes in the ganglion of KAN RNU nude rats to lack T cell markers CD4 or CD8, and this was indeed the case. Despite this, lymphocyte densities in the spiral ganglion were similar in KAN RNU hets and KAN RNU nudes. We asked, what is the identity of these lymphocytes populating the KAN RNU nude ganglia? We found that RNU nude rats have an increased splenic NK cell compartment relative to RNU hets (Figure 5C), consistent with previous reports (Mehrotra et al., 2017; Reynolds et al., 1982). Similarly, in the spiral ganglion, we found lymphocytes positive for the NK cell marker CD161 (Figure 5D). A representative example of the flow cytometry gating strategy used for RNU hets and RNU nude rats is shown in Supplementary Figure 1. In summary, the number of lymphocytes in RNU nude rats is similar to that in RNU het controls in spite of the absence of T cells, likely attributable to an increase in the number of NK cells.

**Figure 5.**
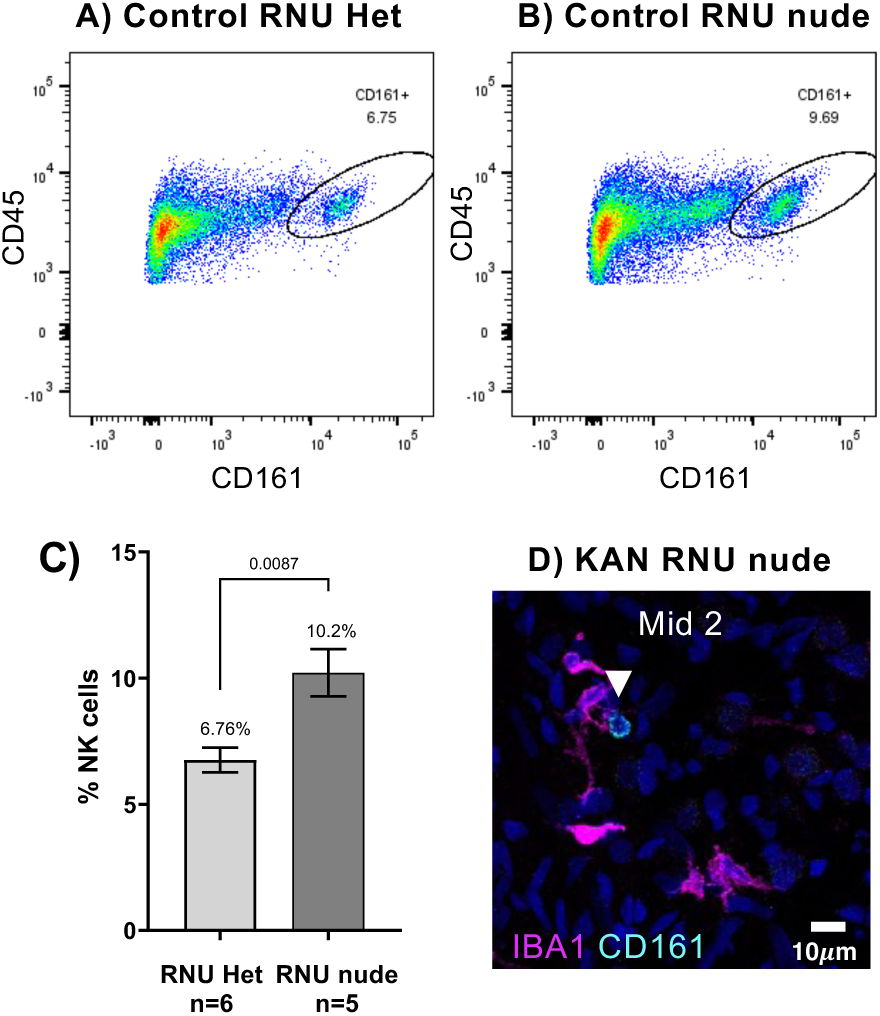
RNU nude rats have increased splenic NK cells. Representative plots from flow cytometric analysis of the spleen showing CD161+ NK cells in control **(A)** RNU het and **(B)** RNU nude rats. CD161+ cells were gated from CD45+/CD3-cells. Upstream gating is shown in Figure 1, and the full gating strategy is shown in Supplementary Figure 1. **(C)** Comparisons of the splenic NK cell compartment indicate a significant increase in the number of NK cells in RNU nude rats compared to RNU hets. Shown are means ± SEM. n=number of animals. Significance was determined by unpaired t-tests. High magnification image of the Mid 2 region of the spiral ganglion in (**D**) labeled to show macrophages (IBA1, magenta), CD161+ NK cells (cyan), and nuclei (blue) in the ganglion of a KAN RNU nude rat. A NK cell is indicated by the white arrow.

### Lymphocytes are depleted in Rag2^-/-^, Il2rg^-/-^ (SRG) rats

SGN death in KAN RNU nude rats occurred at a rate similar to KAN RNU hets, indicating that removing T cells does not prevent or reduce post-deafening SGN death. However, other lymphocytes, particularly NK cells, remain. To assess a possible role for these cells, we evaluated SGN survival in a more severe immunodeficient rat strain, the SRG rat, to assess whether lymphocytes other than T cells may be causal to SGN death after hair cell loss. The SRG (Sprague-Dawley/Rag2^−/−^/Il2rg^−/−^) rat is a severely immunodeficient inbred rat lacking both the Rag2 and Il2rγ genes, and is deficient in mature T, B, and NK cells (Noto et al., 2020). We confirmed lack of lymphocytes in the spleen by flow cytometry (Figure 6A-D). Splenocytes were analyzed for cell size and granularity, and cells in the expected range for lymphocytes were virtually absent in SRG rats, but present in wildtype (WT) controls (Figure 6A-D).

**Figure 6.**
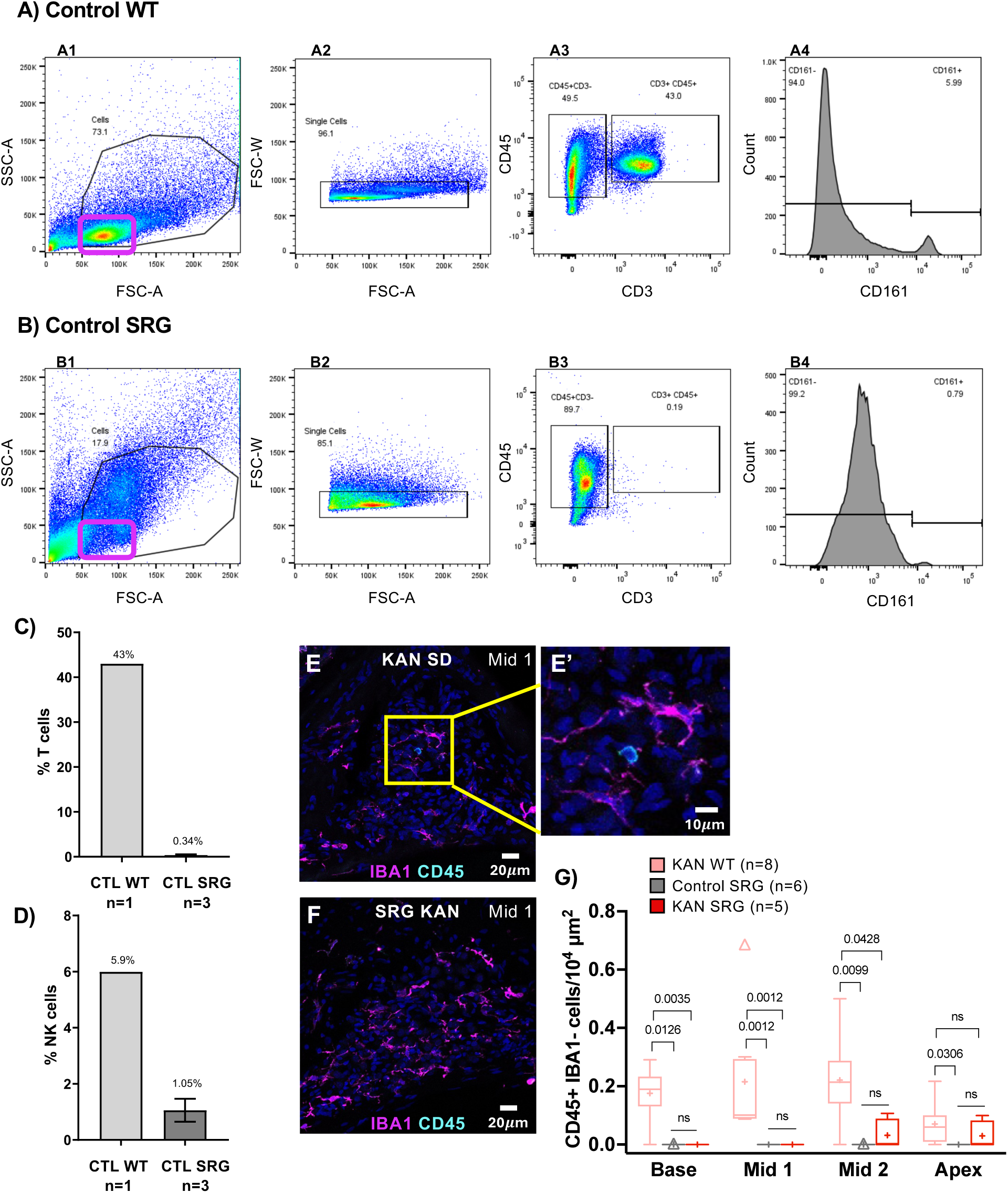
Sprague-Dawley Rag2^-/-^, Il2rg^-/-^ SRG rats are deficient of T, B, and NK cells. Representative plots from flow cytometric analysis of spleen from undeafened **(A)** WT Sprague-Dawley and **(B)** SRG rats. Panels show the gating strategy used to confirm depletion of lymphocytes in the SRG rats. The magenta box in **A1** outlines the population of lymphocytes in WT and **(B1)** lack of cells in this region in the SRG rats. Cells were gated as described in Figure 1. Comparisons between spleens of WT and SRG rats indicate that **(C)** CD3+ T cells and **(D)** CD161+ NK cells are depleted in SRG rats. Shown are means ± SEM. n=number of animals. **(E)** Representative low magnification image of the Mid 1 spiral ganglion region from a KAN WT rat labeled to show the presence of a CD45+/IBA1-cell with characteristic lymphocyte morphology. Yellow box highlights the area shown in **(E’)** at higher magnification. **(F)** CD45+/IBA1-cells are rarely seen in KAN SRG rats using an identical labeling strategy. **(G)** Quantification of CD45+/IBA1-cells in the ganglion of control and KAN SRG rats. Box and whisker plot with box enclosing 25th-75th percentiles, whisker lengths calculated by Tukey’s method, and outliers indicated by individual symbols. Within each box, the horizontal line indicates the median value for each group and the + sign indicates the mean value. Significance of differences was calculated by was determined by Kruskal-Wallis test with Dunn’s correction for multiple comparisons and is indicated by horizontal black bars above the groups. The number of individual animals, n, is in the figure legend.

Having verified lymphocyte depletion in the spleen, we next asked whether lymphocytes persisted in the ganglion of KAN SRG rats. In WT Sprague-Dawley rats and in RNU rats, there was a significant increase in CD45+/IBA-cells post-deafening (Figure 4D, 6E, G). In KAN SRG rats, we found that the spiral ganglion was nearly devoid of CD45+/IBA1-immune cells (Figure 6F, G) – which were completely undetected throughout the ganglion except for 0.2 cells/10^4^ *μ*m^2^ in the mid 2 and apex regions. That CD45+/IBA1-cells are largely absent in SRG rats supports a conclusion that the CD45+/IBA1-immune cells appearing in the deafened ganglia of WT immunocompetent rats are predominately lymphocytes.

### Ablation of T, B, and NK cells in SRG rats results in regional differences in SGN survival after hair cell loss

We next asked whether the absence of lymphocytes in SRG rats affected SGN survival after deafening. Representative images are shown in Figure 7, with quantitation shown in Figure 8A. First, we compared SGN numbers in undeafened control WT Sprague-Dawley and SRG rats to determine whether the Rag2-/-Il2rg-/-mutations affect normal SGN numbers. In the comparison, we found approximately the same number of SGNs throughout the spiral ganglion, except in the apical turn in which SGN density in the SRG rats was slightly but significantly less than in WT controls (Figure 8A).

**Figure 7.**
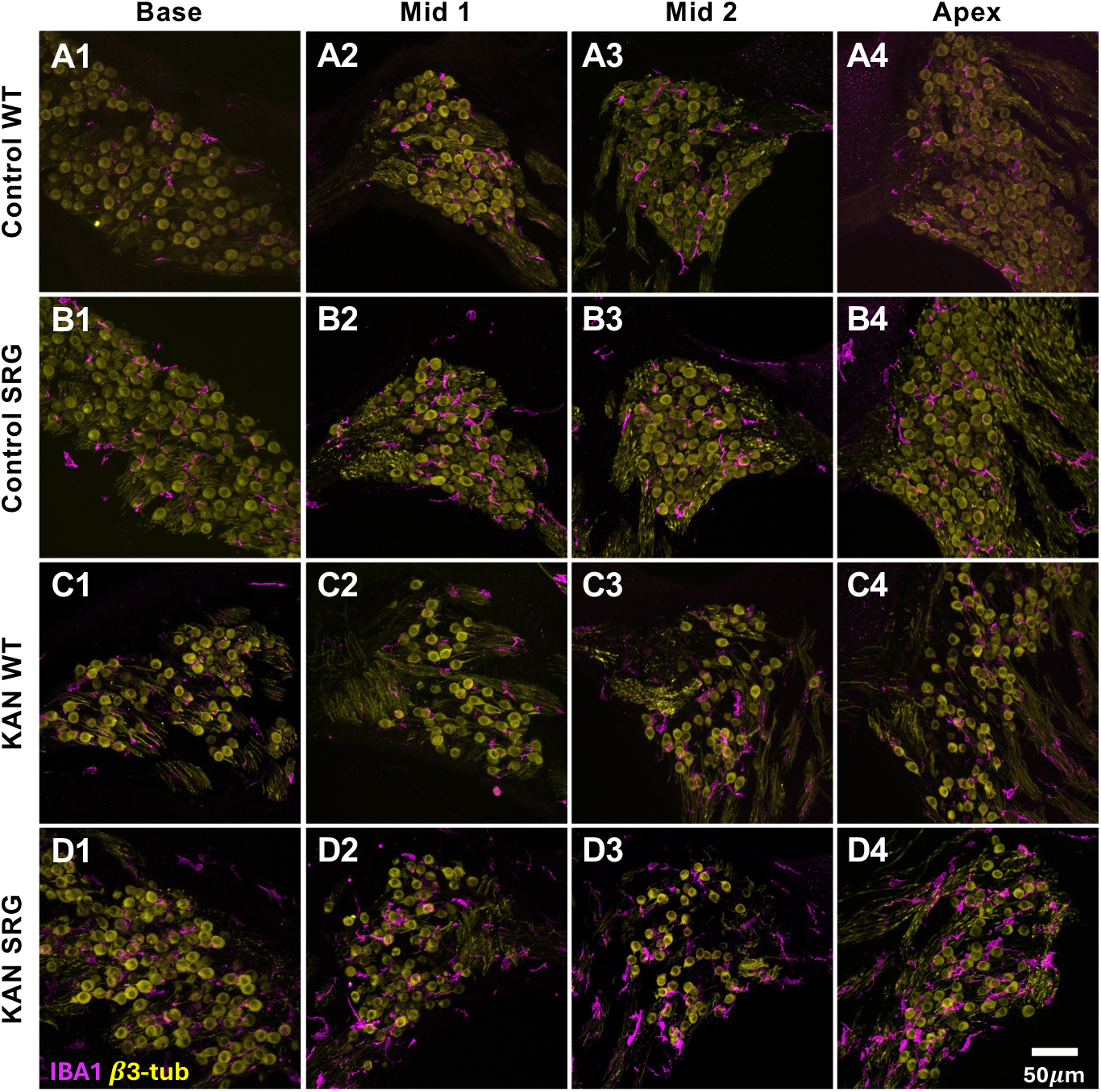
SGNs and macrophages in control and KAN wildtype and SRG rats. Shown are SGNs (βIII-tubulin, yellow) and macrophages (IBA1, magenta) in control wildtype (WT, **A1-A4**), control SRG (**B1-B4**), kanamycin treated (KAN) WT (**C1-C4**), and KAN SRG rats (**D1-4**) at postnatal day 70 (P70). Representative images from cochlear cross-sections at four different locations in the spiral ganglion along the base-to-apex axis (tonotopic) axis are in columns from left to right: column 1 corresponds to the base (column 1), mid 1 (column 2), mid 2 (column 3), and apex (column 4).

**Figure 8.**
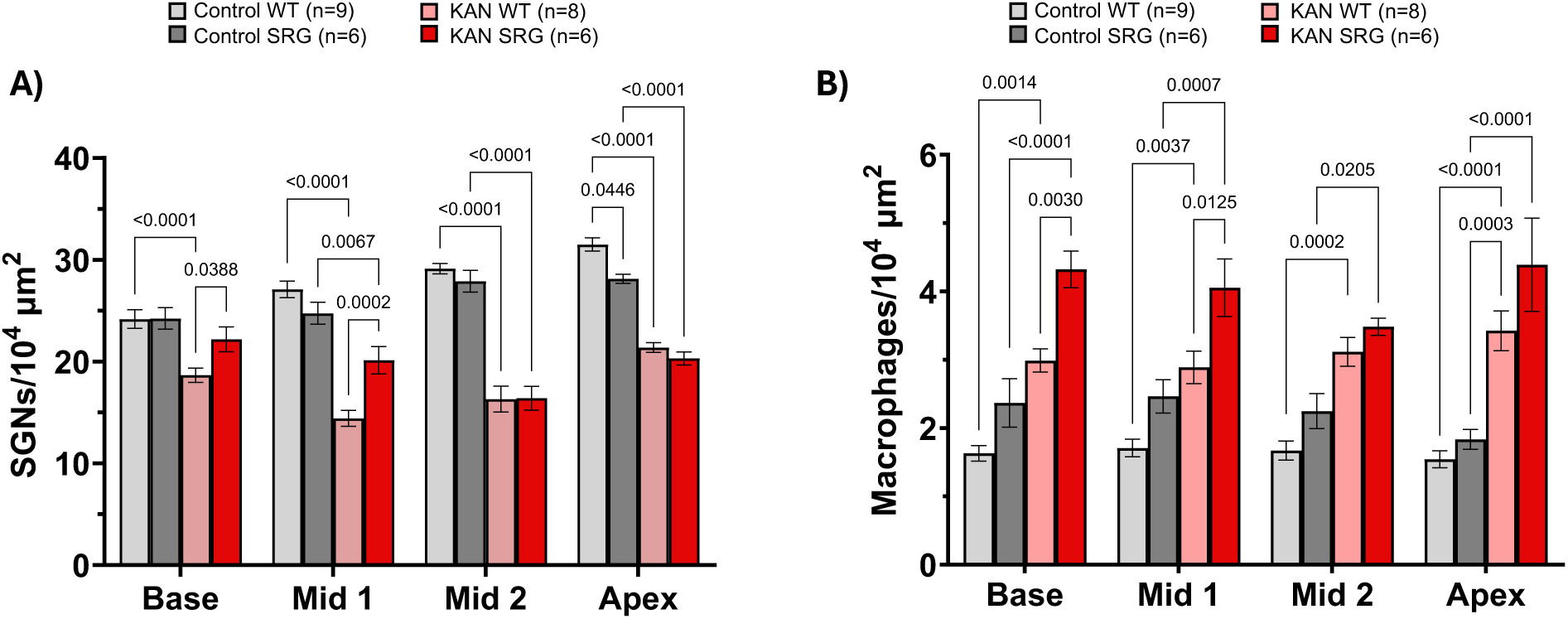
Quantification of SGNs and macrophages by cochlear location in control and KAN SRG rats at P70 indicate regional differences in SGN survival after ablation of T, B, and NK cells. Number of (**A**) SGNs (β_III_-tubulin+ cells) and (**B**) macrophages (IBA1+ cells) per 10^4^ µm^2^ at four locations in the spiral ganglion. Shown are means ± SEM. n=number of animals. Significance of differences was determined using a two-way ANOVA with Tukey’s correction for multiple comparisons. Additionally, because macrophage numbers were greater in all four locations in the spiral ganglion, even if not significantly greater in any individual region, we applied a mixed-effects ANOVA, described in the text in Results, and found that the number of macrophages was overall significantly greater in SRG rats than in WT rats for both control and KAN conditions.

In KAN rats, we found regional differences in SGN loss between SRG and WT rats (Figure 8A). In the basal half – base and mid1 regions – SGN loss post-deafening was significantly reduced in KAN SRG rats relative to KAN WT rats (Figure 8A). Specifically, in the base, there was no apparent neuronal death post-deafening, i.e., no significant difference in SGN density between undeafened control SRG and KAN SRG rats. In the mid 1 region, SGN density was reduced in KAN SRGs relative to undeafened control SRGs but remained significantly greater than in KAN WT rats. These data imply that in the basal half of the spiral ganglion, lymphocytes play a necessary role in SGN death post-deafening. In contrast, in the apical half – mid 2 and apex – this did not appear to be the case; that is, there was no significant difference in SGN density post-deafening between KAN WT and KAN SRG rats. In the mid 2 turn, SGN density was similar between KAN WT and KAN SRG rats, both of which were decreased by approximately 42% relative to respective undeafened controls. In the apex, the density of surviving SGNs was similar between KAN WT and KAN SRG rats but, because the initial number of SGNs was smaller in SRG rats, the magnitude of the decrease was less in KAN SRG rats than in KAN WT rats. These data indicate that lymphocyte deficiency – although not T cell deficiency, as shown above – reduces SGN death, but in a region-dependent manner, specifically in the basal, but not apical, region of the spiral ganglion.

### Spiral ganglion macrophage number increases in KAN WT and KAN SRG rats after hair cell loss

As in the RNU experiments, we asked whether depletion of all lymphocytes in SRG rats affected macrophage recruitment into the spiral ganglion. Representative images are shown in Figure 7, with quantitation in Figure 8B. We first compared macrophage numbers between undeafened control WT Sprague-Dawley and SRG rats. We found a significant increase in spiral ganglion macrophage number in undeafened control SRG rats relative to control WT rats throughout the spiral ganglion. In control WT rats, there were approximately 1.6 cells/10^4^ *μ*m^2^ across all regions of the ganglion, compared to 2.8-3.4 cells/10^4^ *μ*m^2^ in control SRG rats (Figure 8B). A two-way mixed-effects ANOVA (Tukey correction for multiple comparisons) revealed a significant main effect of genotype (F_1,50_ =21, p<0.0001) in control undeafened rats, indicating an overall increase in macrophage number in the SRG rat spiral ganglia.

We have previously shown a post-deafening increase in macrophage number in the ganglia of Sprague Dawley rats (Caro et al., 2026), and we show here that this is similarly the case in KAN SRG rats, particularly in the basal half of the ganglion (Figure 8B). That is, depletion of lymphocytes does not prevent the post-deafening increase in macrophage number. Just as the number of macrophages was larger in undeafened control SRG rats than in control WT rats, the number of macrophages was larger in KAN SRG rats than in KAN WT rats (Figure 8B). For the deafened rats, the two-way mixed-effects ANOVA (Tukey correction for multiple comparisons) revealed a significant main effect of genotype (F_1,50_ =18, p=0.0001). Thus, both control and KAN SRG rats have increased macrophage numbers in the spiral ganglion compared to WT control and KAN rats, respectively, and deafening results in a significant increase in macrophage number for both genotypes.

We have previously shown that the increase in macrophage number post-deafening is coupled with an increase in macrophage activation (Caro et al., 2026; Gansemer et al., 2024), evidenced by presence of CD68 immunoreactivity (Chistiakov et al., 2017; Holness et al., 1993). Here, we evaluated CD68 immunoreactivity in KAN WT and KAN SRG rats to determine macrophage activation status. We found a significant increase in CD68+/IBA1+ double positive cells in KAN SRG rats compared to control SRGs, indicating an increase in the proportion of activated macrophages in the ganglion after deafening (Figure 9A, C, D). Only 2-8% of spiral ganglion macrophages were CD68+ in control SRG rats, whereas in KAN SRGs, 37-47% of macrophages were CD68+, similar to the fraction in KAN WT rats (Figure 9B, D). Few CD68+ macrophages are detected in the ganglia of undeafened WT Sprague-Dawley rats (Caro et al., 2026), and this was similarly the case in the spiral ganglia of undeafened SRGs (Figure 9D). Thus, lymphocytes are not necessary for recruitment nor for activation of macrophages in the spiral ganglion post-deafening.

**Figure 9.**
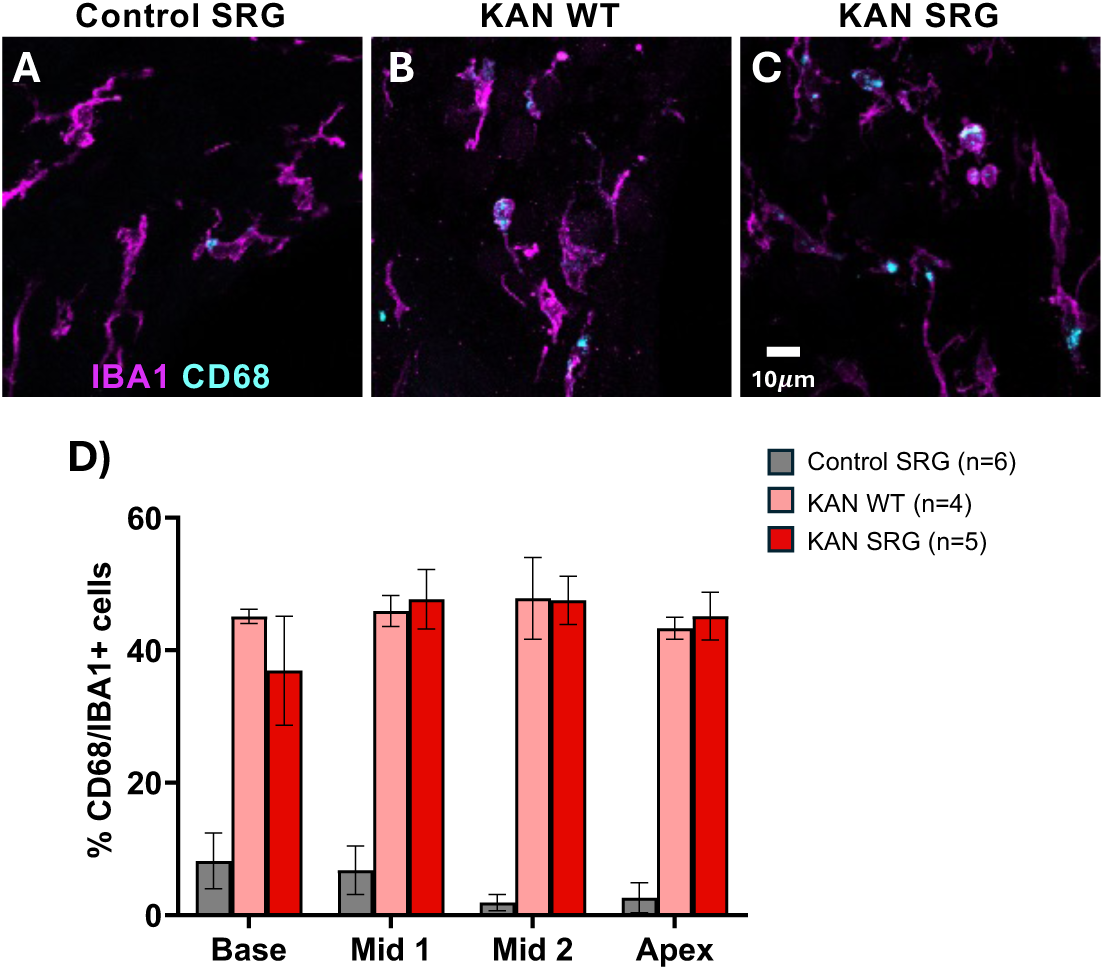
Macrophage activation in the spiral ganglion increases in KAN WT and KAN SRG rats after hair cell loss. Representative high magnification images from the Mid 2 region of spiral ganglia from (**A**) control SRG, (**B**) KAN WT and (**C**) KAN SRG rats. Images labeled to show macrophages (IBA1, magenta) and CD68 (cyan). The percentage of IBA1-positive cells that are also CD68 positive (double positive cells) is shown in **D**. Shown are means ± SEM. n=number of animals. Significance of differences was determined by Mann-Whitney multiple unpaired t-tests.

### Auditory threshold is not affected by Foxn1 or Rag2/Il2rg mutations

We next evaluated whether the Foxn1 or Rag2/Il2rg mutations had any effect on auditory function. We recorded ABR measurements to assess ABR thresholds, wave I amplitudes, and wave I latencies.

Recordings from RNU nude rats were taken between P50-P60 at 4, 8, and 16 kHz. For all frequencies, hearing thresholds were similar for male (Figure 10A) and female (Figure 10B) rats of both genotypes, i.e., RNU het and RNU nude rats. There was a small, but significant increase in wave I amplitude in RNU nude males at 16 kHz and at all frequencies tested for RNU nude females compared to sex-matched RNU hets (Supplementary figure 2A, BC). Wave I latency was longer for female RNU nude rats relative to female RNU hets only at 16 kHz (Supplementary figure 3B).

**Figure 10.**
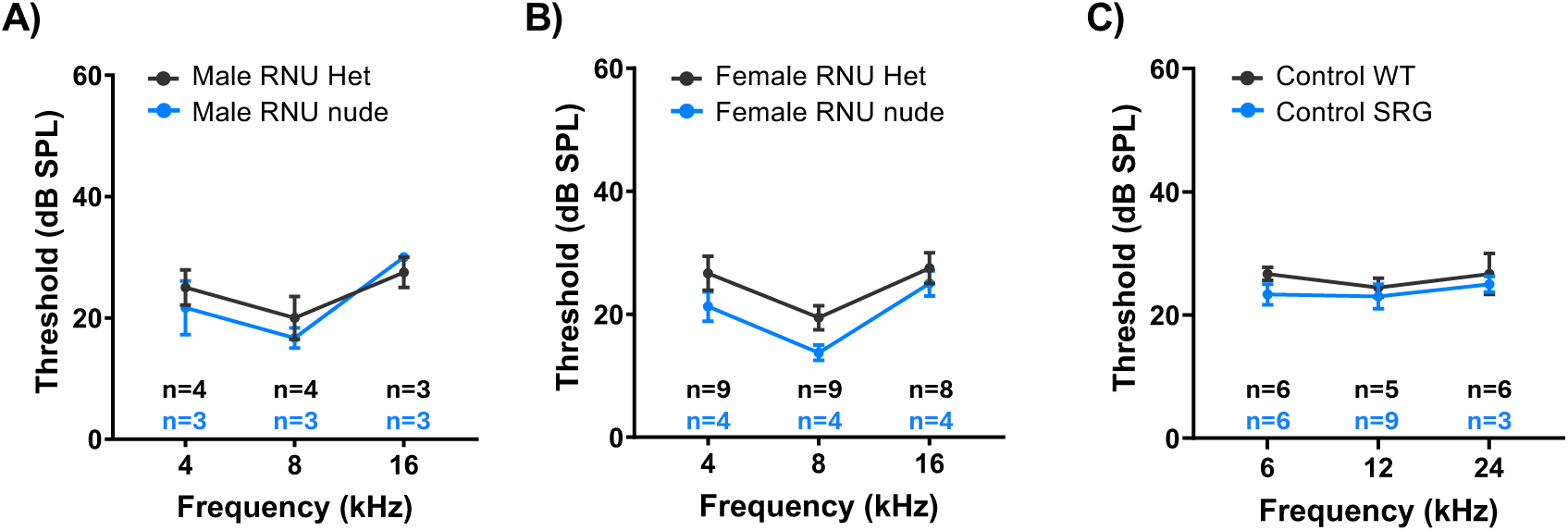
Auditory thresholds are not affected by the Foxn1^rnu/rnu^ or Rag2^-/-^, Il2rg^-/-^ mutations. ABR thresholds for 4, 8, and 16 kHz tone pips in (**A**) male and (**B**) female RNU het and RNU nude rats recorded between P55-P65. n=number of cochlea recorded. **(C)** Thresholds for SRG rats were recorded for 6, 12, and 24 kHz tone pips recorded at P69. n=number of cochlea recorded at each frequency.

Recordings from SRG rats were taken on P69 at 6, 12, and 24 kHz. Recordings for SRGs were from male rats, while male and female WT rats were combined for analysis. ABR thresholds were similar for control WT and control SRG rats at all three frequencies (Figure 10C). Wave I amplitude in control SRG rats was slightly, but significantly, increased at 12 kHz relative to WT (Supplementary figure 4A). Wave I latency was longer in control SRGs relative to WT only at 24 kHz (Supplementary figure 4B).

## Discussion

We have previously shown that the number of macrophages, including MHCII expressing macrophages, increases in the spiral ganglion prior to the onset of post-deafening SGN death (Caro et al., 2026). Additionally, lymphocytes, including helper T cells, cytotoxic T cells, and NK cells, increase in number in the ganglion concurrent with the onset of significant SGN death (Caro et al., 2026). The presence of these cell types indicates the capacity for MHCII mediated antigen presentation, specifically by macrophages to helper T cells, in the ganglion after hair cell loss. Recognition of antigen by T cells results in the activation of T and B lymphocytes, initiating adaptive immune responses. Moreover, helper T cells are orchestrators of the immune response, secreting cytokines to enhance function of immune cells, including NK cells and other lymphocytes, as well as macrophages (S. Chen et al., 2023; Reel et al., 2024).

Upregulation of MHCII molecules and T cell infiltration into the brain has been linked to neuron loss in CNS neurodegenerative conditions such as Parkinson’s (Brochard et al., 2009; Subbarayan et al., 2020) and Alzheimer’s diseases (X. Chen et al., 2023; Zeng et al., 2024). Reports have shown that in the absence of T cells, neuron loss in the substantia nigra is reduced in a rat model of Parkinson’s disease (Subbarayan et al., 2020) and hippocampal volume is maintained in an Alzheimer’s tauopathy model (X. Chen et al., 2023), implying a causal role for T cells in neurodegeneration. Given the similar increase in the number of CD4+ T cells and MHCII+ macrophages in the cochlea post-deafening or after noise-exposure (Yang et al., 2015), a similar role for antigen-presentation in cochlear neurodegeneration may exist. Here, we used two immunodeficient rat strains to investigate the role of lymphocytes and the adaptive immune response in SGN death post-deafening.

To directly test the specific requirement for T cells in SGN death post-deafening, we used RNU nude rats that entirely lack T cells. These were compared to RNU heterozygote littermates that have a normal T cell complement. We found that across all regions of the ganglion, the extent of SGN loss was similar in KAN Het and KAN RNU rats, indicating that T cell responses are not necessary for post-deafening SGN death (Figure 2, 3A) nor are they necessary for the post-deafening increase in macrophages within the spiral ganglion (Figure 2, 3B). However, NK cells persist in the RNU nude ganglia in greater numbers than in RNU hets and increase in number post-deafening (Figures 4, 5). These observations that both spiral ganglion macrophages and NK cells increase in number post-deafening despite the absence of T cells imply that T cells are not required for innate immune responses in the cochlea.

NK cells are lymphocytes of the innate immune response that, upon activation, mediate cellular cytotoxicity through the release of perforin and granzymes, leading to the death of target cells. To assess whether NK cells play a role in spiral ganglion neurodegeneration post-deafening, we evaluated SGN survival in the T, B, and NK cell deficient SRG rat. The results suggest a role for NK cells, but to a differing extent along the tonotopic axis of the spiral ganglion. Deletion of lymphocytes in the SRG rat had a significant effect on the extent of SGN death (Figure 7) and extent of macrophage recruitment (Figure 8) in the basal half of the ganglion, but not the apical half. We infer that the effect of the SRG gene mutations on reducing SGN death is attributable, specifically, to lack of NK cells because lack of T cells in RNU rats had no such effect, ruling out significant involvement of the adaptive immune system. Moreover, we have not observed B cells in the spiral ganglion in any circumstance.

These results raise the question of why NK cells should be playing a greater role in neurodegeneration in the basal half of the ganglion than the apical half. One potential explanation arises from previous gene expression profiling analysis which shows differential expression of several cytokines post-deafening (Rahman et al., 2023). Among these are IL-18 and IL-1β, which are both members of the IL-1 family of pro-inflammatory cytokines (Sims & Smith, 2010). Increased expression of IL-1β has been observed in several CNS neurodegenerative diseases, including multiple sclerosis and Alzheimer’s disease (Griffin et al., 1989; Patterson, 1995). IL-1β is produced by both neurons and immune cells (Guo et al., 2003) and its receptor, IL-1R1 is similarly expressed by neurons and immune cells (Copray et al., 2001), allowing for both paracrine and autocrine signaling. Activation of the IL-1R1 complex initiates a signaling cascade that drives the expression of cytokines (Garlanda et al., 2025) and recruitment and activation of immune cells. In particular, IL-18 is a potent activator of NK cells (Hyodo et al., 1999; Srivastava et al., 2010; Tsutsui et al., 2003), and the pro-inflammatory cytokine IL-1β synergizes with IL-18 in activation of NK cells (Dinarello, 2018). Notably, both IL-18 and IL-1β are expressed at a significantly higher levels in the basal half of the ganglion post-deafening, as is a gene encoding the IL-1 receptor complex member IL1RAP (Rahman et al., 2023). A higher level of activating factors, such as IL-18 and IL-1β, and a cognate receptor could result in a higher level of NK cell activity and consequent cytotoxicity in the basal half of the ganglion, accounting for the significantly better SGN survival when NK cells are removed. Conversely, in the apical half of the ganglion, less activation of NK cells would diminish their role in SGN death and reduce the effect of their removal.

These results further imply that neurodegeneration post-deafening in the apical half of the spiral ganglion has a different cellular basis than in the basal half. In neither region does depletion of T cells affect neurodegeneration. Additionally depleting NK cells does reduce neurodegeneration in the basal, but not in the apical half. Macrophages are implicated in neurodegeneration post-deafening, as depletion of macrophages reduces SGN death in kanamycin-deafened mice (Shimada et al., 2023), and we suggest a greater role for macrophages in the apical half of the spiral ganglion. Through much of the period of SGN death, which is at a similar rate throughout the spiral ganglion in deafened wildtype rats, macrophages are more abundant in the apical than in the basal spiral ganglion (Caro et al., 2026), as are levels of macrophage chemoattractants such as CCL2 and CCL7 (Rahman et al., 2023). A possibly greater role for macrophages in the apical half could account for the equivalence of SGN death in the apical and basal halves, despite possibly greater NK cell activity in the basal half.

## Acknowledgements

We thank Drs. Marlan Hansen, Noah Butler, Dan Summers, and Michael Dailey for valuable comments on the manuscript, Catherine Kane for maintaining our animal colony and technical assistance, and Tristan Brown for assistance in labeling of cochlear sections. This work is based on research available as a preprint from BioRxiv https://dx.doi.org/10.64898/2026.03.10.710901. This work was supported by the National Institute on Deafness and other Communication Disorders grants F31 DC021590 (AMC) and R01 DC015790 (SHG). The data presented herein were obtained in part at the University of Iowa Carver College of Medicine/Holden Comprehensive Cancer Center Flow Cytometry core research facility, which is funded through user fees and the generous financial support of the Carver College of Medicine, Holden Comprehensive Cancer Center, and Iowa City Veteran’s Administration Medical Center.

## Supplementary Figures

**Supplementary figure 1.**
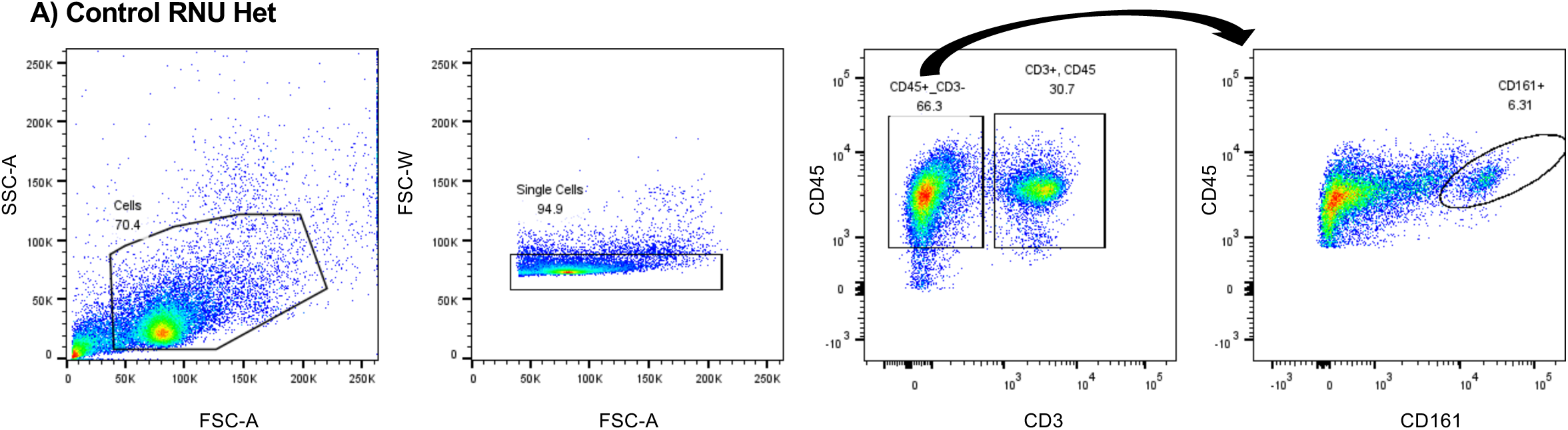
Flow cytometry gating strategy. Representative plots from flow cytometric analysis of spleen from a RNU heterozygous rat. As in Figure 1, panels show first total cells captured. The first gate was set from the forward-scatter versus side-scatter to capture cells in the expected size and granularity range for lymphocytes. Single cells were then selected by size; doublet cells were excluded. The cells captured within the single cell gate were then plotted to evaluate expression of CD45 and CD3. CD45+/CD3-cells were then evaluated for CD161 expression, as noted by the black arrow. Labels inside each plot indicate the target population captured within the gate. The numbers in each plot indicate the percent of cells represented within the gate.

**Supplementary figure 2.**
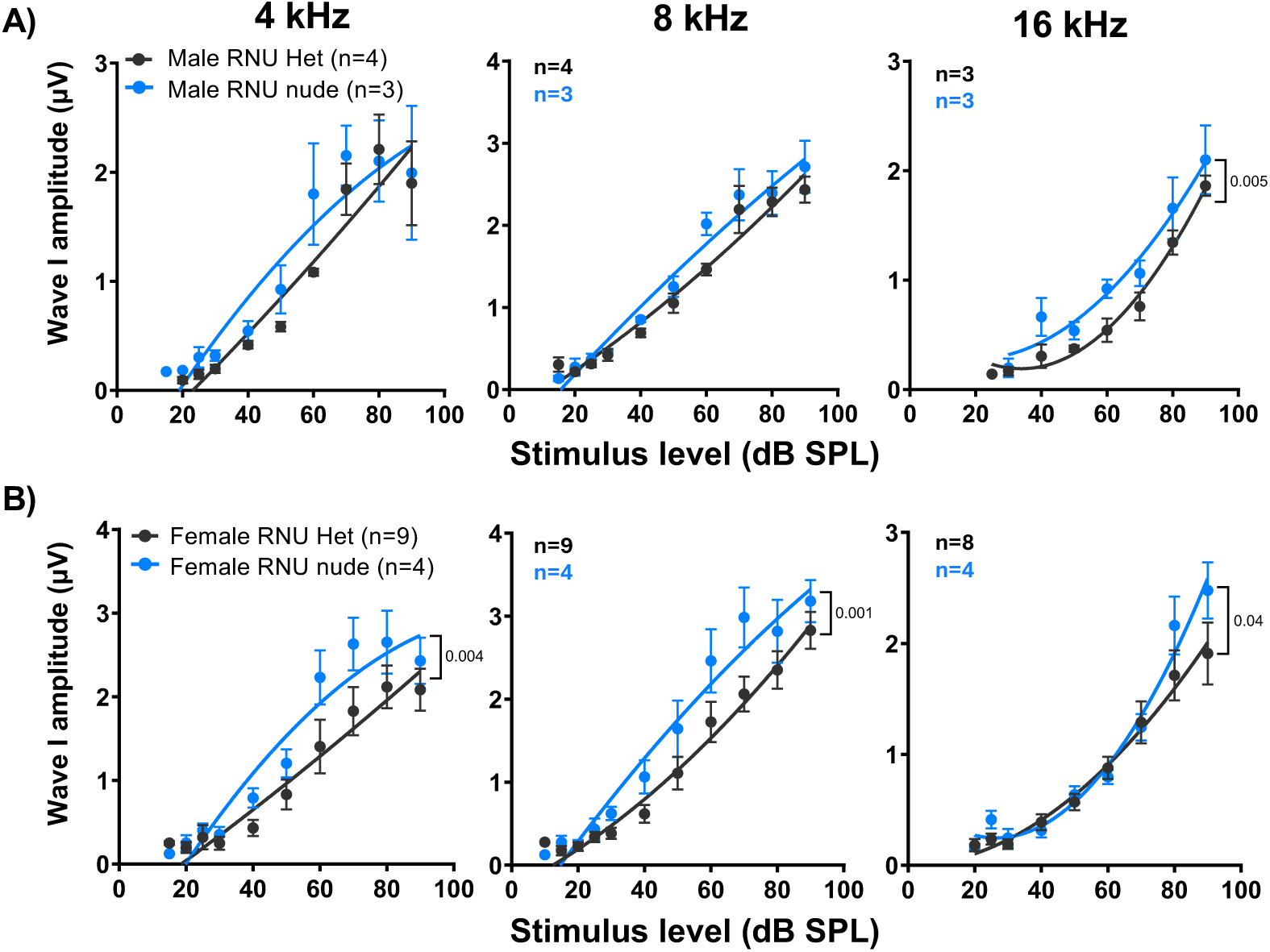
Wave I amplitudes for male and female RNU rats. As shown in Figure 10, ABR thresholds were recorded for 4, 8, and 16 kHz tone pips. Shown here are wave I amplitudes for (A) male and (B) female RNU rats. Data were fit to a second order polynomial and significance of differences for each plot was determined by an extra sum-of-squares F test. Number of cochlea recorded is indicated for each frequency.

**Supplementary figure 3.**
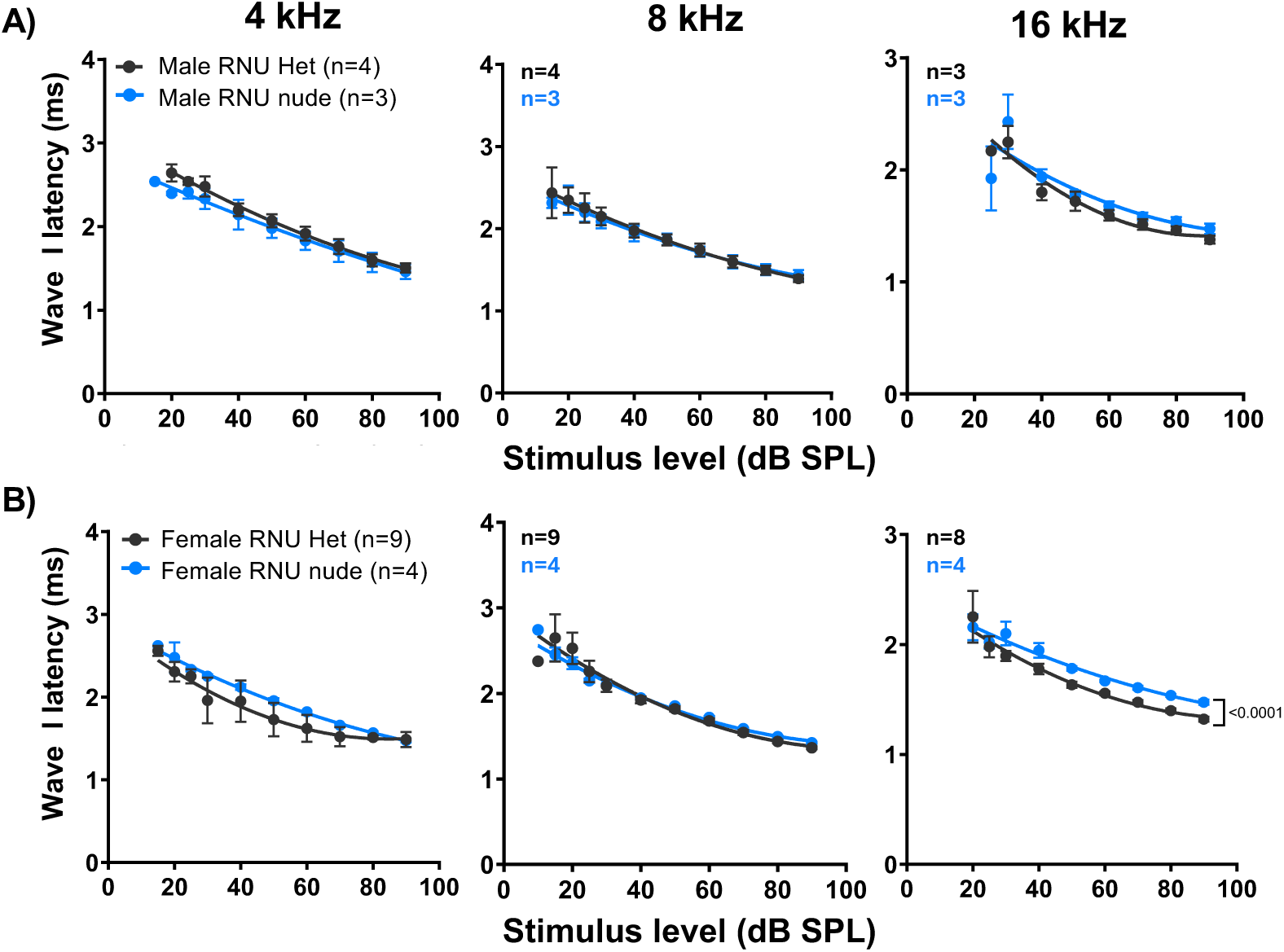
Wave I latencies for male and female RNU rats. As shown in Figure 10, ABR thresholds were recorded for 4, 8, and 16 kHz tone pips. Shown here are wave I latencies for (A) male and (B) female RNU rats. Data were fit to a second order polynomial and significance of differences for each plot was determined by an extra sum-of-squares F test. Number of cochlea recorded is indicated for each frequency.

**Supplementary figure 4.**
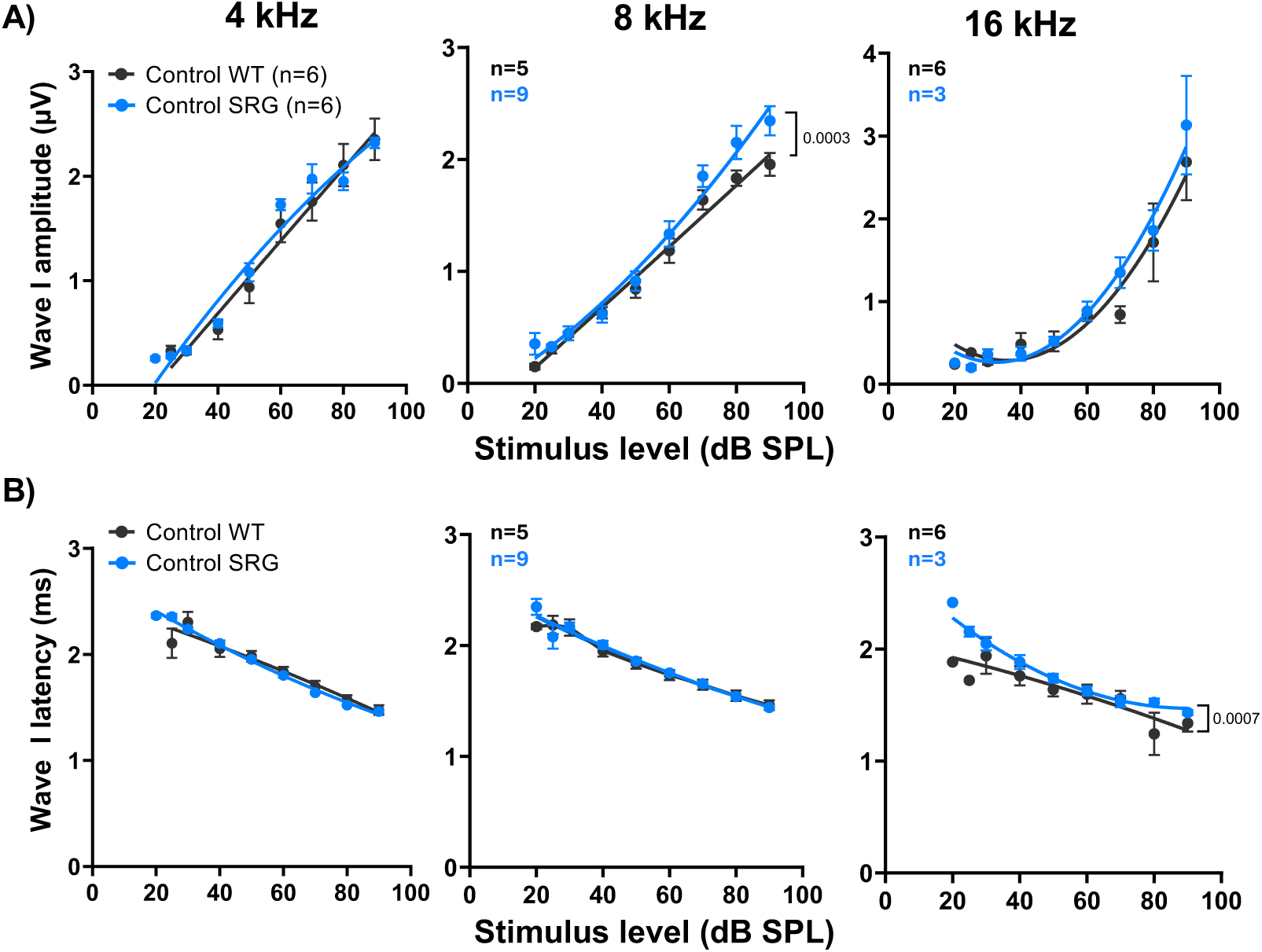
Wave I amplitudes and wave I latencies for SRG rats. **(A)** ABR thresholds for 6, 12, and 24 kHz tone pips in WT and SRG rats recorded at P69. Recordings from male and female WT rats were combined, but, because of limited availability, only male SRG rats were tested. Graphs showing **(A)** wave I amplitude and **(B)** wave I latency are shown for each frequency. Data were fit to a second order polynomial and significance of differences for each plot was determined by an extra sum-of-squares F test. Number of cochlea recorded is indicated for each frequency.

## Notes

https://github.com/bgansemer/ABR-analysis

### Competing Interest Statement

The authors have declared no competing interest.

### Summary of Updates

Figure 6 has been reformatted to fit on a single page and some of the accompanying text was revised.

